# The predicted RNA-binding protein ETR-1/CELF1 acts in muscles to regulate neuroblast migration in *Caenorhabditis elegans*

**DOI:** 10.1101/2020.02.25.965277

**Authors:** Matthew E. Ochs, Matthew P. Josephson, Erik A. Lundquist

## Abstract

Neuroblast migration is a critical aspect of nervous system development (e.g., neural crest migration). In an unbiased forward genetic screen, we identified a novel player in neuroblast migration, the ETR-1/CELF1 RNA binding protein. CELF1 RNA binding proteins are involved in multiple aspects of RNA processing including alternative splicing, stability, and translation. We find that a specific mutation in alternatively-spliced exon 8 results in migration defects of the AQR and PQR neurons, and not the embryonic lethality and body wall muscle defects of complete knockdown of the locus. Surprisingly, ETR-1 was required in body wall muscle cells for AQR/PQR migration (i.e. it acts cell non-autonomously). Genetic interactions indicate that ETR-1 acts with Wnt signaling, either in the Wnt pathway or in a parallel pathway. Possibly, ETR-1 is involved in the production of a Wnt signal or a parallel signal by the body wall muscles that controls AQR and PQR neuronal migration. In humans, CELF1 is involved in a number of neuromuscular disorders. If the role of ETR-1/CELF1 is conserved, these disorders might also involve cell or neuronal migration. Finally, we describe a technique of amplicon sequencing to detect rare, cell-specific genome edits by CRISPR/Cas9 *in vivo* (CRISPR-seq) as an alternative to the T7E1 assay.

## Introduction

The migration of neuroblasts and neurons during development is a tightly orchestrated process that is imperative for the proper development and function of the nervous system. To understand the fundamental mechanisms used in the migration of neurons and neural crest neuroblasts, we utilize the simple model organism nematode worm *Caenorhabditis elegans*. The Q neuroblasts in *C. elegans* are bilaterally symmetrical neuroblasts that are an ideal model for directed cell migration and discovery of genetic mechanisms that control directed cell migration undergo similar division patterns as well as migration patterns (reviewed in (Middelkoop and Korswagen 2014)) (Chapman *et al*. 2008; Sundararajan and Lundquist 2012; Josephson *et al*. 2016). The stereotyped simplicity of Q neuroblast migration involves two distinct phases (Figure 1) (Sulston and Horvitz 1977; Chalfie and Sulston 1981; Chapman *et al*. 2008; Sundararajan and Lundquist 2012). QR and QL are born in between the hypodermal seam cells V4 and V5 in the posterior-lateral region of the animal, QR on the right and QL on the left. In the first phase, QR migrates anteriorly over the V4 seam cell, and QL posteriorly over the V5 seam cell, at which point the first cell division occurs. The second phase involves a series of migrations, divisions, and cell death resulting in the production of three neurons: AQR, AVM, and SDQR from QR; and PQR, AVM, and SDQL. QR descendants migrate anteriorly, and QL descendants posteriorly. The first phase of migration is controlled by the transmembrane receptors UNC-40/DCC, PTP-3/LAR, and MIG-21 (Middelkoop *et al*. 2012; Sundararajan and Lundquist 2012), and also involves the Fat-like cadherins CDH-3 and CDH-4 (Sundararajan *et al*. 2014; Ebbing *et al*. 2019). The second phase of long-range Q descendant migration is controlled by Wnt signaling, in both canonical and non-canonical roles (see (Eisenmann 2005; Zinovyeva *et al*. 2008; Ji *et al*. 2013; Josephson *et al*. 2016)).

**Figure 1.**
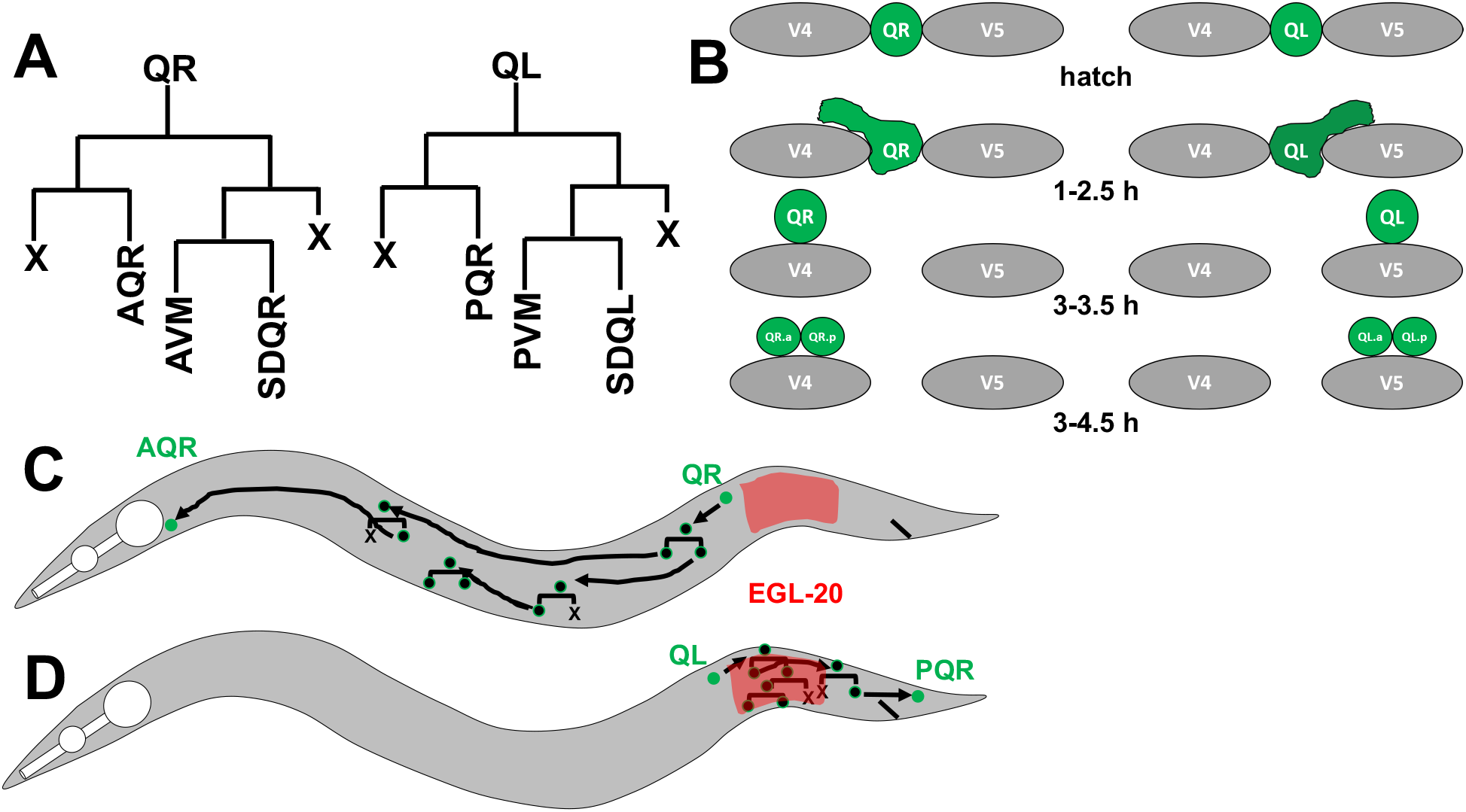
Depiction of Q cell and descendant migration. A) The lineages of QL and QR cell descendants. A programmed cell death is indicated with an “X”. B) Early Q migration. At hatching, QL and QR are rounded and unpolarized. At 1-2.5 h after hatching, QR extend a cellular protrusion to the anterior over the V4 seam cell, and QL extends a protrusion to the posterior over V5. At 3-3.5 h after hatching, the cell body migrates along the path of the earlier protrusion, and the first cell division occurs after migration at 3-4.5 h post hatching. C) A depiction of QR migration. QR migrates anteriorly and does not respond to the EGL-10/Wnt signal (red), and Q descendants migrate anteriorly. D) QL migrates posteriorly and responds to the EGL-20/Wnt signal via canonical Wnt signaling and BAR-1/ß-catenin. This activates the expression of the MAB-5/Hox transcription factor in QL, which is necessary and sufficient to drive continued posterior Q descendant migration.

A forward genetic screen for defects in migration of the Q descendants AQR and PQR identified the *etr-1(lq61)* mutation. ETR-1 is an ELAV-type RNA-binding protein similar to mammalian CELF-1 (CUGBP ELAV-like family member 1) implicated in numerous neurodegenerative and neuromuscular disorder including myotonic dystrophy type I (Li *et al*. 2001; Savkur *et al*. 2001; Timchenko *et al*. 2001; Timchenko *et al*. 2004; Ho *et al*. 2005; Kuyumcu-Martinez *et al*. 2007; Sofola *et al*. 2007; Sofola *et al*. 2008; Daughters *et al*. 2009; Schoser and Timchenko 2010; Wijsman *et al*. 2011; Berger and Ladd 2012). CELF1 molecules have been shown to regulate multiple aspects of mRNA processing including translational regulation, mRNA stability, and alternative splicing (reviewed in (Dasgupta and Ladd 2012)). Previous work has shown that RNAi knockdown of *etr-1* results in severe muscle disorganization and the Paralyzed arrest at two-fold elongation (Pat) phenotype (Milne and Hodgkin 1999). Accordingly, *etr-1* was expressed in the body wall musculature (Milne and Hodgkin 1999), consistent with the role of human CELF1 in myotonic dystrophy type I. recent studies show that ETR1 is expressed in all cells in the embryo (Boateng *et al*. 2017). Thus, ETR-1/CELF1 molecules are evolutionarily-conserved regulators of muscle development and physiology. Recent work has shown that depletion of *etr-1* results in germline defects including smaller oocytes, reduced fertility, and a failure to engulf germ cells undergoing programmed cell death (Boateng *et al*. 2017), a non-muscle role of ETR-1. While *etr-1* RNAi resulted in Pat animals and embryonic lethality, the *etr-1(lq61)* allele described here is viable and fertile.

In this work we describe a role of ETR-1 in Q neuroblast migration. *etr-1* mutants resulted in misplaced AQR and PQR, but did not affect early Q migration, suggesting a defect in phase II but not phase I of migration. The *etr-1* locus is extensively alternatively spliced (WormBase web site, http://www.wormbase.org, release WS274, 9/27/2019)(Boateng *et al*. 2017), similar to CELF1 family members in other species (Li *et al*. 2001; Barreau *et al*. 2006). The viable *etr-1* mutation was a premature stop codon in alternatively-spliced exon 8, suggesting that a loss of a subset of *etr-1* transcripts containing exon 8 perturbed cell migration but did not result in lethality. Cell-specific expression experiments and cell-specific CRISPR/Cas9 genome editing indicated that ETR-1 was required in body wall muscles and not in the Q cells themselves for AQR and PQR migration, a non-autonomous role. Finally, *etr-1* interacted genetically with *Wnt* mutations in a manner consistent with ETR-1 affecting Wnt signaling. Our results suggest that *etr-1* mutation pertubs muscle differentiation, one consequence of which is to disrupt a muscle-derived guidance signal for AQR and PQR, possibly a Wnt signal or a parallel pathway.

CELF1 has been associated with multiple neurodegenerative and neuromuscular disorders, including Myotonic Dystrophy type I (DMI) (Savkur *et al*. 2001; Timchenko *et al*. 2001; Timchenko *et al*. 2004; Ho *et al*. 2005; Kuyumcu-Martinez *et al*. 2007; Schoser and Timchenko 2010; Berger and Ladd 2012), the cardiac syndrome arrythmogenic right ventricular dysplasia (Li *et al*. 2001) and neurological disorders such as Alzheimer’s disease (Wijsman *et al*. 2011), spinocerebellar ataxia type 8 and possibly fragile X syndrome (Sofola *et al*. 2007; Daughters *et al*. 2009). The role of ETR-1 in body wall muscles (Milne and Hodgkin 1999) is consistent with the role of CELF1 in neuromuscular disease is humans. If other roles are conserved, cell or neuroblast migration might be a conserved component of one or more of these human disorders.

## Materials and Methods

### Data and reagent Availability

All data, reagents, and strains used in this work are freely available upon request.

### Strains and Genetics

*C. elegans* were grown using standard methods at 20°C. N2 Bristol strain was used as wildtype. Alleles used include LGI: *lin-44(n1792), mab-5(e1239)*, LGII: *etr-1(lq61 and lq133), cwn-2(ok564)*, LG IV: *egl-20(n585), egl-20(gk453010), egl-20(mu39), cwn-2(ok895)*. Standard gonad injection was used to create extrachromosomal arrays: *lqEx912, lqEx913, and lqEx914 [Pegl-17::etr-1::GFP* (25 ng/μL), *Pscm::GFP* (25 ng/μL)]; *lqEx944, lqEx945, lqEx946, lqEx947 [Pmyo-3::etr-1* (25 ng/μL), *Pscm::GFP* (25 ng/μL). For these cell-specific rescue experiment, the entire *etr-1* coding region, from initiator methionine to the stop codon in the full-length isoform, was amplified by PCR, including introns, and placed behind the *egl-17* and *myo-3* promoters. The coding regions of *etr-1* were sequenced to ensure no errors were introduced as the result of PCR.

### Cell-specific somatic CRISPR/Cas9 genome editing

We used the cell-specific CEISPR protocols as previously described (Shen *et al*. 2014), involving expression of Cas9 from cell-specific promoters and ubiquitous expression the th sgRNA from the ubiquitous small RNA U6 promoter. Ultraviolet trimethylpsoralen (UV/TMP) techniques were used to integrate the following extrachromosomal arrays into transgenes: LGII: *lqIs244 [Pgcy-32::CFP* (25 ng/μL)], unknown chromosomal location *lqIs327 [Pmyo-3::Cas9/etr-1 sgRNA* (25 ng/μL), *Pgcy-32::YFP* (25 ng/μL)], *lqIs308(Pegl-17::Cas9/mab-5* sgRNA (25 ng/μL), *Pgcy-32::YFP* (25 ng/μL)]. The sequences of all plasmids and primers used to construct them are available upon request.

### AQR/PQR forward genetic mutant screen and mapping

Standard techniques were used mutagenize L4 and young adult hermaphrodites harboring a *Pgcy-32::cfp* (strain LE2500) with ethylmethane sulfonate (EMS) using standard techniques (Anderson 1995). Mutagenized animals were plated on single seeded NGM plates and allowed to self-fertilize. F1 animals were picked to plates, with three animals per plate. F2 progeny were screened with a fluorescence dissecting microscope, and animals with misplaced AQR and/or PQR visualized by the *lqIs58[Pgcy-32::cfp]* transgene were isolated. Germline mutation was confirmed by screening the progeny for AQR and PQR defects. Approximately 3000 haploid genomes were screened. New mutations were mapped using single nucleotide polymorphism mapping combined with next generation sequencing using the Cloudmap pipeline and the polymorphic CB4856 Hawaiian strain (Davis *et al*. 2005; Minevich *et al*. 2012).

### Scoring Q-cell and AQR/PQR migration defects

To score early Q cell migrations we expressed GFP using the seam cell promoter *(Pscm)* expressed in the hypodermal seam cells and the Q cells. as described previously (Chapman *et al*. 2008; Dyer *et al*. 2010; Sundararajan and Lundquist 2012). Briefly, adults were allowed to lay eggs overnight and adults and larvae were washed away with M9 buffer, leaving the eggs on the plate. Larvae were collected every hour via M9 buffer washing, and aged to the appropriate development stage (2.5-4.5 h post hatching) before imaging. At least 50 cells were scored for each genotype and statistical significance was determined using Fisher’s Exact Test.

Previously-described techniques were used to quantify AQR and PQR migration (Chapman *et al*. 2008; Sundararajan and Lundquist 2012). AQR migrates to and resides just behind the posterior pharyngeal bulb in the anterior deirid ganglion, and PQR migrates to and resides just behind the anus in the phasmid ganglion. We used *Pgcy-32::CFP* to visualize AQR and PQR. We used five positions along the anterior-posterior axis to score AQR and PQR (Figure 2C). Position 1 is the wildtype location of AQR and is around the posterior pharyngeal bulb. Position 2 was posterior of position 1, but anterior to the vulva. Position 3 is the region around the vulva. Position 4 is where the Q cells are born. Position 5 is the wildtype location of PQR and is just posterior to the anus. Fisher’s exact test was used to test for significance. The predicted additive phenotype of double mutants was calculated by the formula p(A) = p1 + p2 – (p1p2), where p(A) is the predicted additive proportion, p1 is the proportion in single mutant 1, and p2 is the proportion in single mutant 2.

**Figure 2.**
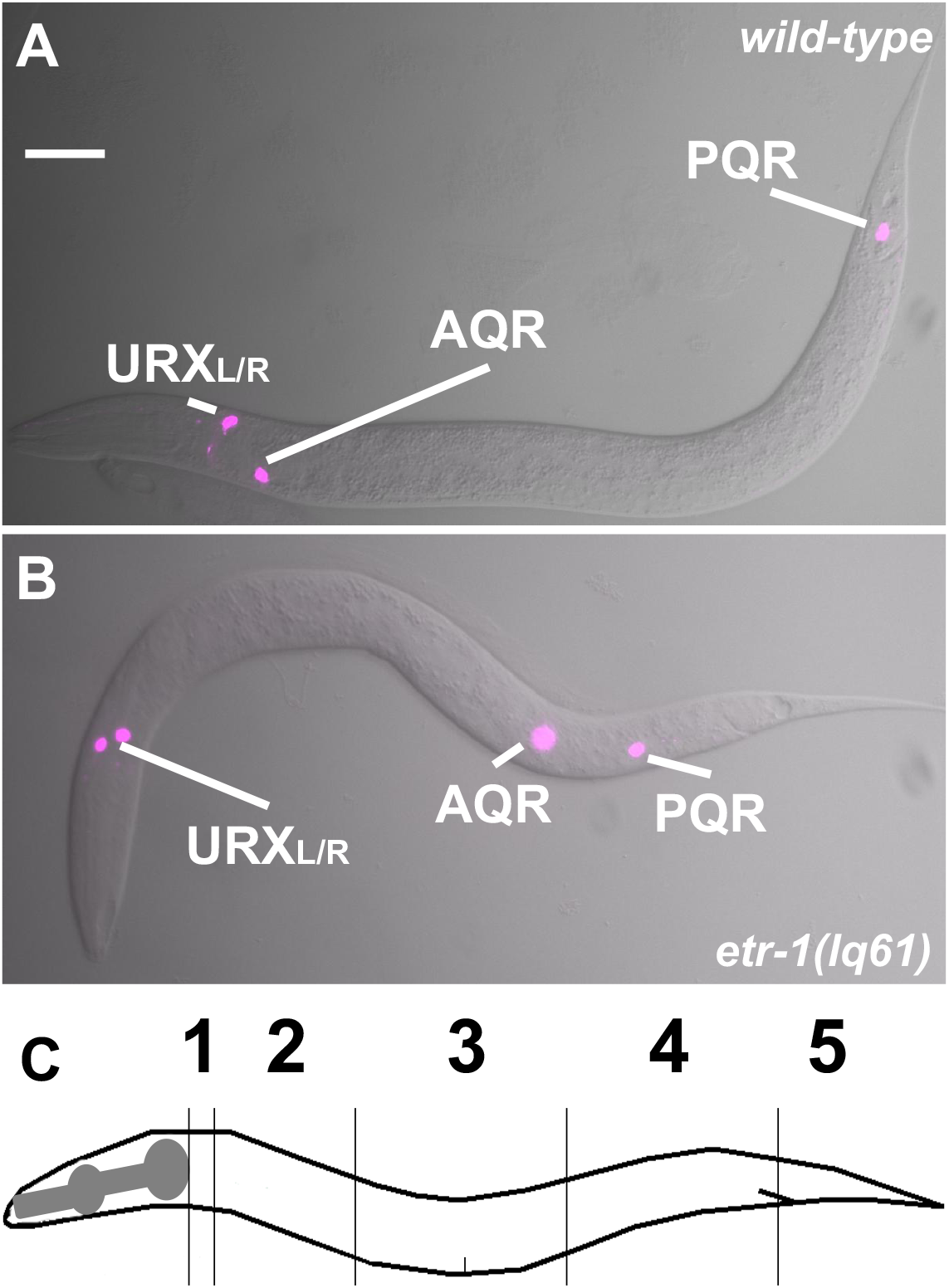
*etr-1(lq61)* causes AQR and PQR migration defects. A and B are merged Differential Interference Contrast and *cyan fluorescent protein* images of AQR and PQR via the *gcy-32::cfp* transgene *lqIs58*, also expressed in the URX neurons. A) An L4 wild-type animal with AQR and PQR in their normal positions, near the posterior pharyngeal bulb and posterior to the anus, respectively. B) An *etr-1(lq61)* L4 animals displayed AQR and PQR migration defects. C) A depiction of the scoring positions of AQR and PQR in Tables 1–3. Position 1 is the normal position of AQR, and position 5 the normal position of PQR.

### CRISPR-seq: whole-genome sequencing to detect cell-specific genome edits

Amplicon sequencing of the *etr-1* exon 8 region surrounding the sgRNA site was conducted using a two-step PCR protocol to amplify the genomic region and to attach Illumina-specific clustering and sequencing adaptors. For PCR round 1 (PCR1), forward and reverse primers were designed to amplify a 170-bp region surrounding the *etr-1* sgRNA site, 24 bp from the PAM site. The “smRNA” Illumina adapter primer was added to the 3’ end of the forward gene-specific sequence, and the Illumina “Read2” primer was added to the 3’ end on the reverse complementary gene-specific primer. PCR1F and PCR1R primers (15 pMol each) were used in 15 cycles of PCR (20ul reaction) on 50ng of genomic DNA harvested from wild-type N2 and from animals harboring an integrated transgene expressing Cas9 and the sgRNA *(lqIs327* for *etr-1*, and *lqIs308* for *mab-5*).

### PCR1 (20ul)

3μl DNA (50ng)

1μl forward primer (gene specific) 15pMol

1 μl reverse primer (gene specific) 15 pMol

5μl ddH20

10μl PCR master mix

### PCR1 cycling

94° 2 min

94° 10 sec

60° 30 sec

68° 1 min

goto step 2 14x

For PCR round 2 (PCR2), primers used were a forward Illumina i5 dual index sequencing primer TagAAACGG and a unique i7 reverse primers (*e.g*. i7_i07 (CAGATC) for N2 DNA, and i07_i08 (ACTTGA) for *etr-1* muscle-specific CRISPR). The i5 and i7 primers (15pMol each) were used for 10 cycles of amplification of 15μl of 1:10 diluted PCR1 (50μl reaction volume). The i5 and i7 primers have sequence overlap with the smRNA and Read2 sequences used in PCR1 and become incorporated into the PCR2 product.

### PCR2 (50μl)

15μl diluted PCR1 DNA

2μl i5 primer 15 pMol

2μl i7 primer (gene specific index) 15 pMol

6μl ddH20

25μl PCR master mix

### PCR2 cycling

94° 2 min

94° 10 sec

60° 30 sec

68° 1 min

goto step 2 9x

Agencourt Ampure beads were used to remove salt, enzyme, and unincorporated nucleotides and primers. 200μl of Agencourt Ampure beads were mixed with with 25μl of PCR2 in 75μl of elution buffer (5mM Tris-HCl pH 8.5) for 100μl total. PCR products were allowed to bind to the beads for 10 minutes. Tubes were placed on a magnetic bead stand to clear the beads to the side of the tube, and allowed to sit for 5 minutes. Supernatant was carefully removed with a 100μl pipet tip. Beads were washed with 50μl of freshly-prepared 80% Ethanol in ddH_2_0 for 1 minute. Supernatant was removed carefully, and another wash with 80% Ethanol was performed. After removing supernatant, the beads were spun in a microcentrifuge at full speed for 30 seconds. Remaining Ethanol was carefully removed, and the beads were allowed to air dry for 5 minutes. PCR products were eluted by adding 17μl of 10mM Tric-HCl pH 8.5 with 0.05% Tween-20. After mixing and a 5-minute incubation, the tube was placed back on the magnet stand for 5 minutes, and the eluate was carefully removed and placed in a fresh tube.

One μl of eluate was analyzed on a Tapestation fragment analyzer. For *etr-1* exon 8, an amplicon of 301 bp is expected. Smaller fragments were observed which likely represented primer dimers. As long as there is was substantial amount of amplicon present (~one quarter of the product), the primer dimers did not significantly impinge on read counts upon sequencing.

Sequencing was performed on an Illumina Miseq flow cell using a high percentage (20%) of PhiX174 DNA because of the low complexity of the samples. A minimum of 100,000 reads per sample were generated. If a large amount of primer dimer is detected, a higher read count is required. FastqGroomer (Blankenberg *et al*. 2010) was used for read quality control, and were aligned to the *C. elegans* genome using BWA MEM (Li and Durbin 2009) and the *C. elegans* WS220 ce10 reference genome build using the Galaxy platform (Afgan *et al*. 2018). Alignment BAM files were analyzed with the Integrated Genome Viewer (Robinson *et al*. 2011; Thorvaldsdottir *et al*. 2013; Robinson *et al*. 2017), from which coverage and base deletion information were extracted and plotted in Figure 5. Results were visualized using the R package “ggplot2” (Wickham 2009; Team 2017).

A similar procedure was used to sequence amplicons from similar targeting of the *mab-5* locus with expression of Cas9 from the *egl-17* promoter expressed in the Q neuroblasts. The primers used in this *mab-5* experiment are listed (mab-5PCR1F and mab-5PCR1R).

### Primers

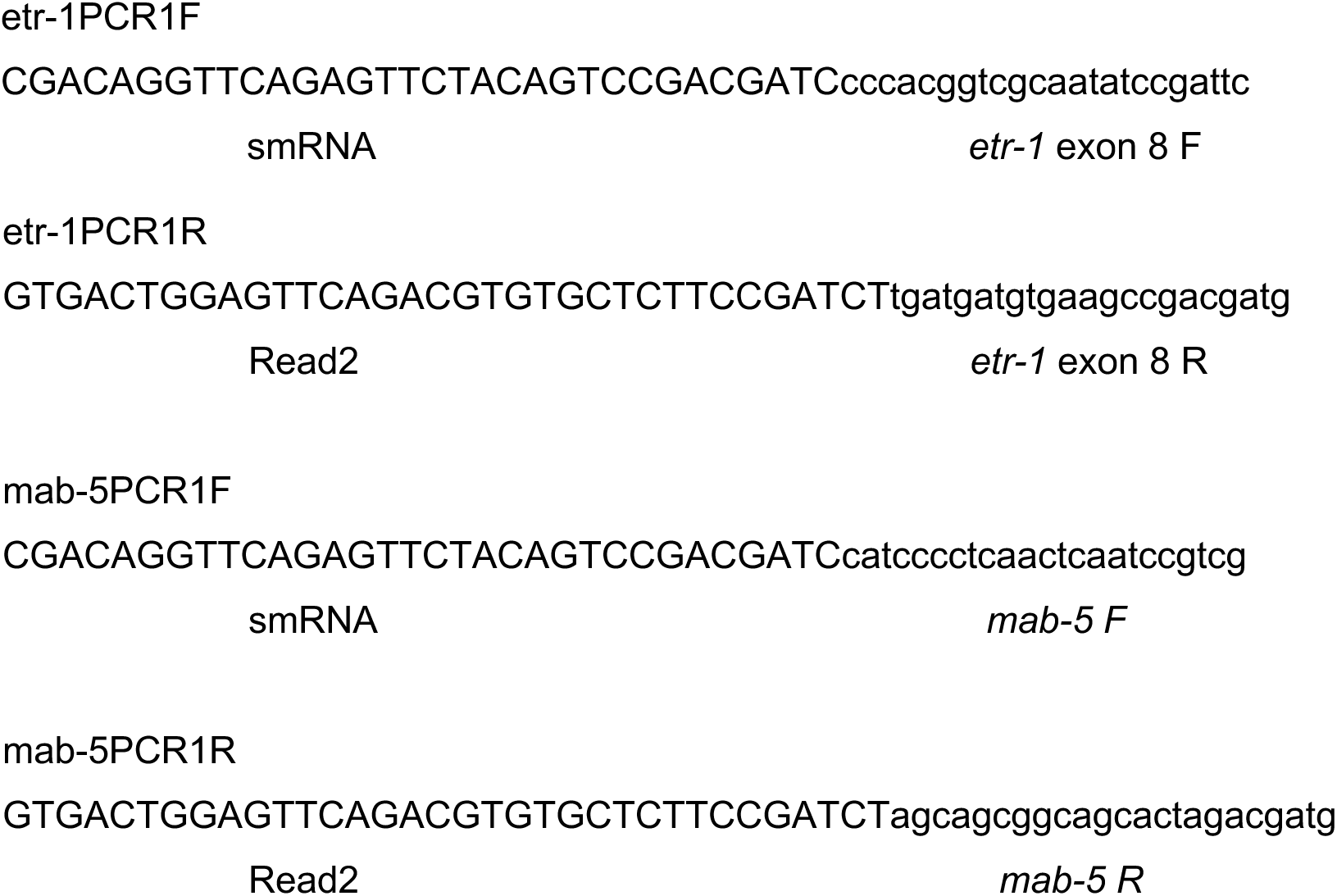

## Results

### Mutations in *etr-1/CELF* perturb directional AQR and PQR migration

The bilateral Q neuroblasts, QL on the left and QR on the right, undergo an identical pattern of cell division and cell death to produce three neurons each (Figure 1A) (Sulston and Horvitz 1977; Chalfie and Sulston 1981; Middelkoop and Korswagen 2014). However, QL and descendants migrate posteriorly, and QR and descendants migrate anteriorly (Figure 1B-D). Initial Q migration at 1-2.5 h after hatching involves protrusion to the posterior and anterior for QL and QR respectively, followed by migration of the cell bodies to positions above the V5 or V4 seam cells (Figure 1B) (Honigberg and Kenyon 2000; Chapman *et al*. 2008). The first Q cell divisions occur between 3-4.5 h after hatching. Initial Q migration is independent of Wnt signaling and involves the transmembrane receptors UNC-40/DCC, PTP-3/LAR, and MIG-21, and the Cadherins CDH-4 and CDH-3 (Honigberg and Kenyon 2000; Middelkoop *et al*. 2012; Sundararajan and Lundquist 2012; Sundararajan *et al*. 2014; Ebbing *et al*. 2019). The second phase of Q descendant migration involves Wnt signaling (Figure 1C,D)(reviewed in (Middelkoop and Korswagen 2014)), (Kenyon 1986; Salser and Kenyon 1992; Chalfie 1993; Harris *et al*. 1996; Whangbo and Kenyon 1999; Korswagen *et al*. 2000; Herman 2001; Eisenmann 2005; Sundararajan *et al*. 2015; Josephson *et al*. 2016). As QL migrates posteriorly, it encounters a posteriorly-derived EGL-20/Wnt signal, which, via canonical Wnt signaling, leads to the expression of MAB-5/Hox in QL and descendants. MAB-5/Hox is both necessary and sufficient for continued posterior Q descendant migration (Figure 1C). QR does not respond to the EGL-20/Wnt signal, does not express MAB-5/Hox, and therefore migrates anteriorly (Figure 1D). Of the Q descendants, PQR migrates furthest to the posterior behind the anus, and AQR migrates furthest anteriorly to a position near the posterior pharyngeal bulb (Figure 1C,D) (Sulston and Horvitz 1977; White *et al*. 1986; Chapman *et al*. 2008). The position of AQR and PQR has been a sensitive method to identify new mutations that perturb Q migrations (Chapman *et al*. 2008; Sundararajan *et al*. 2014; Sundararajan *et al*. 2015; Josephson *et al*. 2017).

A forward genetic screen for new mutations affecting AQR and PQR migrations identified the *lq61* allele. *lq61* mutants displayed weak but significant defects in both the extent and direction of AQR and PQR (Table 1 and Figure 2). The cell migration defects of the *lq61* strain, isolated in the N2 strain background, were mapped relative to single nucleotide polymorphisms (snps) in the CB4856 strain using the Cloudmap sequencing and snp mapping protocol (Minevich *et al*. 2012). This analysis revealed that *lq61* was linked to snps on the far-left end of linkage group II (Figure 3A,B). In this interval was a premature stop codon in exon 8 of the *etr-1* locus, which encodes the *C. elegans* molecule most closely related to mammalian CELF1 (Figure 3B and C) (Dasgupta and Ladd 2012). ETR-1 contains three predicted RNA Recognition Motifs (RRMs) characteristic of the CELF1 family (Figure 3D).

**Figure 3.**
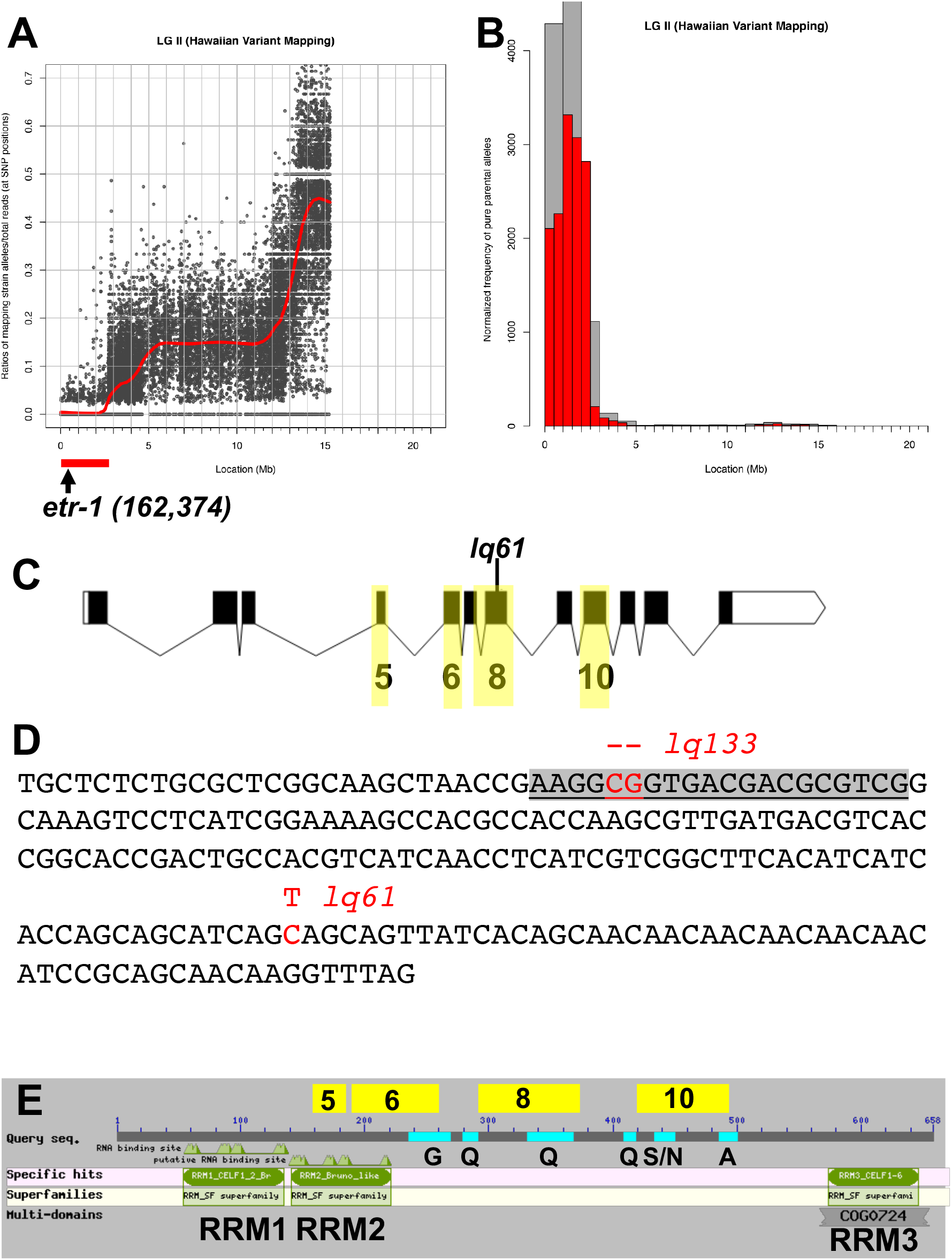
The *etr-1* locus and molecule. A and B) Output file from a Cloudmap experiment showing the relative proportions of HA snps (Y axis) along chromosome II (X-axis) after outcrossing *lq61* to the HA CB4856 strain, and isolating 10 independent mutant lines from the progeny of the heterozygote. These ten lines were combined and subject to whole genome sequencing. The low level of HA snps at the far-left arm of chromosome II indicates the likely position of *lq61*. C) This region contained a premature stop codon in the *etr-1* gene, which starts at nucleotide 162,374 of chromosome II. Boxes represent exons and lines introns. Black boxes represent open reading frame coding region. There is an untranslated exon to the 5’ (exon 1) not included in this depiction. The exons highlighted in yellow are those that display alternative exon usage in *etr-1* isoforms. D) The nucleotide sequence of *etr-1* exon is shown. *lq61* is a C to T transition resulting in a premature stop codon. *lq133* was generated using CRISPR/Cas9 genome editing. The sgRNA sequence is highlighted in grey. *lq133* is a 2-bp deletion at ther PAM site of the sgRNA, and results in a frame shift and premature stop codon. E) The structure of the full-length 658-residue ETR-1 polypeptide. The three RNA Recognition Motifs are indicated, as are the glycine-rich (G), glutamine-rich (Q), alanine-rich (A), and serine/asparagine-rich (S/N) regions. The regions coded for by alternatively-spliced exons are indicated above the line.

**Table 1.** *etr-1* mutant AQR/PQR migration defects.

| Genotype | AQR |  |  |  |  | *p* | PQR |  |  |  |  | *n* | *p* |
| --- | --- | --- | --- | --- | --- | --- | --- | --- | --- | --- | --- | --- | --- |
|  | 1 | 2 | 3 | 4 | 5 |  | 1 | 2 | 3 | 4 | 5 |  |  |
| *wild-type* | 100 | 0 | 0 | 0 | 0 |  | 0 | 0 | 0 | 0 | 100 | 100 |  |
| *etr-1(lq61)* | 85 | 12 | 2 | 0 | 1 |  | 3 | 5 | 4 | 0 | 88 | 100 |  |
| *etr-1(lq133)* | 75 | 19 | 13 | 3 | 0 |  | 1 | 1 | 5 | 10 | 82 | 100 |  |
| *Ex[etr-1(BWMCRISPR)]* | 86 | 13 | 1 | 0 | 0 |  | 1 | 0 | 2 | 1 | 96 | 100 |  |
| *etr-1(lq61); Ex[etr-1(+)] #1* | 97 | 2 | 0 | 0 | 1 | *< 0.01 | 0 | 0 | 0 | 0 | 100 | 100 | *<0.01 |
| *etr-1(lq61); Ex[etr-1(+)] #2* | 98 | 2 | 0 | 0 | 0 | *< 0.01 | 0 | 0 | 0 | 0 | 100 | 100 | *<0.01 |

\*compared to *etr-1(lq61)* at AQR position 1 and PQR position 5.

To confirm that AQR/PQR defects in *lq61* are due to mutation of *etr-1*, we used CRISPR/Cas9 to generate a 2bp frame shift allele *Iq133* in exon 8 (Figure 3C). *etr-1(lq133)* animals also displayed AQR/PQR migration defects similar to *etr-1(lq61)* (Table 1). A fosmid clone containing the wild-type *etr-1(+)* gene rescued *etr-1(lq61)* (Table 1). Together, these results indicate that *etr-1* regulates directed AQR/PQR migration.

Previous studies showed that RNAi of *etr-1* results in embryonic lethality with body wall muscle attachment defects (Milne and Hodgkin 1999). Both *lq61* and *lq133* mutants are viable and fertile, suggesting that they disrupt a subset of ETR-1 functions. The *etr-1* locus is extensively alternatively spliced, including exon skipping of exons 5, 6, 8, and 10 (Figure 3B) (WormBase web site, http://www.wormbase.org, release WS274, 9/27/2019). Indeed, the *lq61* and *Iq133* mutations reside in alternatively-spliced exon 8 of *etr-1*, that is included in only 19 of more than 100 known ETR-1 isoforms (WormBase web site, http://www.wormbase.org, release WS274, 9/27/2019). These results suggest that splice isoforms of *etr-1* containing exon 8 are not required for viability or fertility, but are required for AQR/PQR directional migration. Exon 8 encodes a variable region of CELF-family molecules between RRMs 2 and 3. In ETR-1, this region does not contain recognizable domains, but does include sequences rich in glutamine (Q) residues (Figure 3D). Interestingly, none of the exon 8-containing isoforms also contain exon 10 (WormBase web site, http://www.wormbase.org, release WS274, 9/27/2019), which encodes serine/asparagine-rich and alanine-rich regions (Figure 3D).

### *etr-1* exon 8 mutations do not affect early Q protrusion or migration

In the early L1 after hatching, QL and QR undergo their initial migrations (Figure 1B). QL on the left protrudes and migrates posteriorly over the V5 seam cell, and QR on the right protrudes and migrates anteriorly over the V4 seam cell (Figure 4A and B). Defects in the direction of initial Q protrusion and migration can result migration defects of Q descendants AQR and PQR. *etr-1(lq61)* mutation had no effect on initial QL or QR protrusion or migration: QL protruded and migrated posteriorly, and QR anteriorly (Figure 4C and D) (n = 50). These data suggest that ETR-1 is not required for initial Q protrusion and migration, but rather is specifically involved in Q descendant migration.

**Figure 4.**
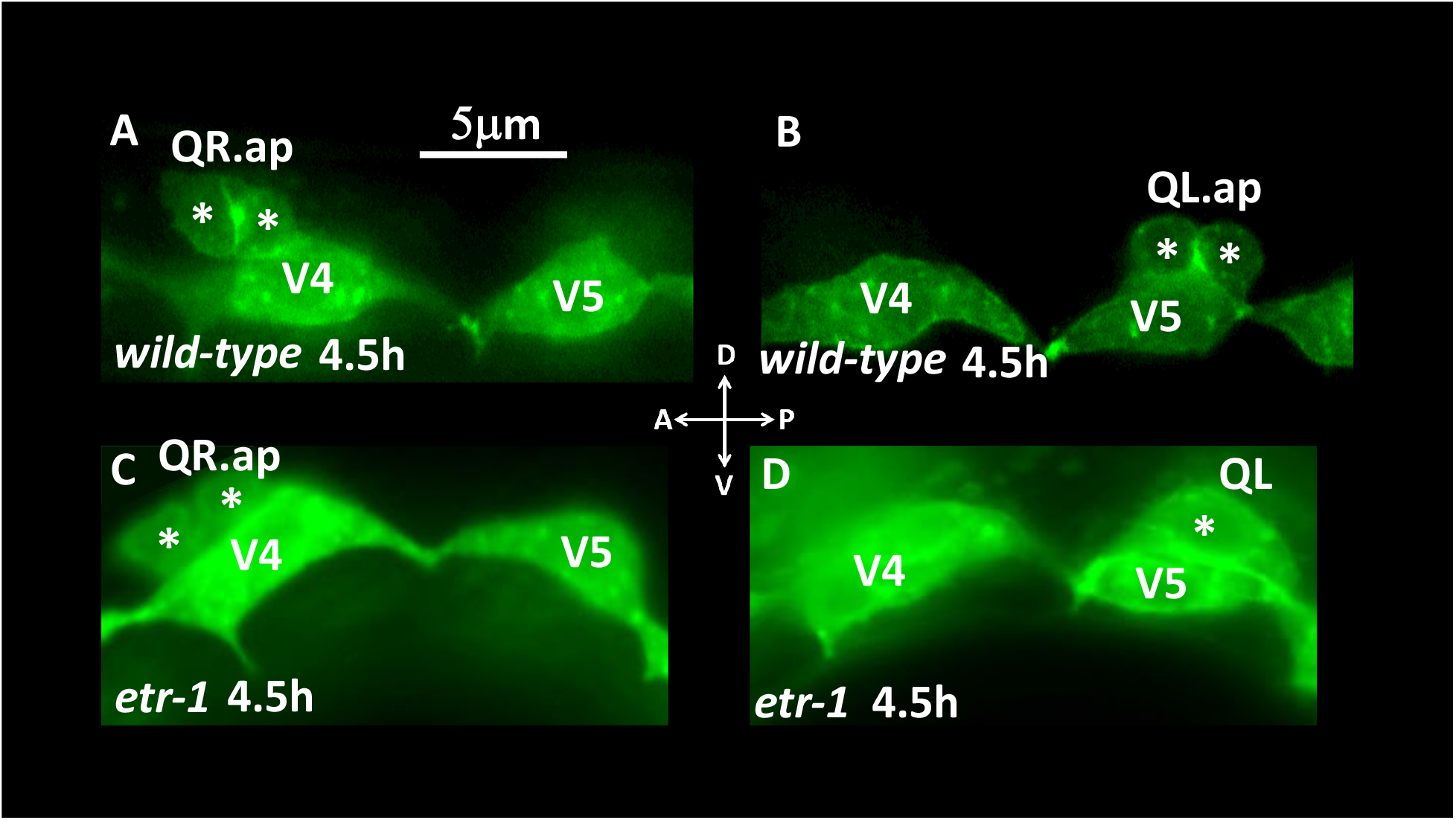
Early Q migrations. Fluorescent micrographs of L1 animals expressing *scm::gfp* transgene *lqIs80* at the indicated time post-hatching are shown. Dorsal is up, and anterior to the left. Asterisks indicate the Q cells and descendants. A) A wild-type QR migrated anteriorly and divided atop the V4 seam cell. B) A wild-type QL migrated posteriorly and divided atop the V5 seam cell. C) An *etr-1(lq61)* mutant QR migrated anteriorly and divided atop the V4 seam cell. D) An *etr-1(lq61)* QL migrated posteriorly atop the V5 seam cell. QR and QL migrated normally in 50 *etr-1(lq61)* animals examined.

### *etr-1* is required in body wall muscle cells for AQR/PQR migration

Previous studies showed that *etr-1* was expressed in body wall muscles and was required for muscle attachment and function (Milne and Hodgkin 1999). To determine where *etr-1* was required for AQR/PQR defects, we used cell-specific expression of the *etr-1(+)* genomic locus, from the initiator methionine to the stop codon, including all introns, driven by cell-specific promoters. Expression of *etr-1(+)* driven from the Q-cell-specific *egl-17* promoter did not rescue AQR and PQR migration defects (Table 2). Expression from the body-wall muscle-specific *myo-3* promoter rescued AQR/PQR defects (Table 2), suggesting that *etr-1* function in muscles is important for AQR/PQR migration.

**Table 2.** *etr-1* rescue by expression in Q cells and body wall muscles.

| Genotype | AQR |  |  |  |  | *p* | PQR |  |  |  |  | *n* | *p* |
| --- | --- | --- | --- | --- | --- | --- | --- | --- | --- | --- | --- | --- | --- |
|  | 1 | 2 | 3 | 4 | 5 |  | 1 | 2 | 3 | 4 | 5 |  |  |
| Q cell expression *Ex[egl-17::etr-1(+)]* |  |  |  |  |  |  |  |  |  |  |  |  |  |
| *etr-1(lq61); Ex #1* | 85 | 11 | 0 | 2 | 1 | NS | 0 | 0 | 0 | 9 | 91 | 100 | NS |
| *etr-1(lq61) no Ex* | 84 | 14 | 0 | 0 | 2 |  | 0 | 0 | 0 | 10 | 90 | 100 |  |
| *etr-1(lq61); Ex #2* | 88 | 7 | 1 | 2 | 2 | NS | 0 | 0 | 0 | 11 | 89 | 100 | NS |
| *etr-1(lq61) no Ex* | 85 | 10 | 1 | 1 | 3 |  | 0 | 0 | 0 | 8 | 92 | 100 |  |
| *etr-1(lq61); Ex #3* | 87 | 10 | 2 | 1 | 0 | NS | 1 | 0 | 5 | 4 | 90 | 100 | NS |
| *etr-1(lq61) no Ex* | 83 | 13 | 2 | 2 | 0 |  | 1 | 1 | 3 | 6 | 89 | 100 |  |
| Body wall muscle expression *Ex[myo-3::etr-1(+)]* |  |  |  |  |  |  |  |  |  |  |  |  |  |
| *etr-1(lq61); Ex #1* | 97 | 0 | 0 | 0 | 0 | < 0.001 | 2 | 0 | 0 | 0 | 95 | 97 | < 0.01 |
| *etr-1(lq61) no Ex* | 70 | 22 | 5 | 1 | 2 |  | 2 | 1 | 3 | 7 | 87 | 100 |  |
| *etr-1(lq61); Ex #2* | 86 | 14 | 0 | 0 | 0 | 0.051 | 0 | 0 | 0 | 2 | 98 | 100 | 0.010 |
| *etr-1(lq61) no Ex* | 74 | 24 | 1 | 1 | 0 |  | 4 | 3 | 0 | 5 | 88 | 100 |  |
| *etr-1(lq61); Ex #3* | 94 | 6 | 0 | 0 | 0 | 0.023 | 0 | 0 | 0 | 1 | 99 | 100 | NS |
| *etr-1(lq61) no Ex* | 81 | 14 | 2 | 1 | 2 |  | 1 | 0 | 0 | 0 | 99 | 100 |  |
| *etr-1(lq61); Ex #3* | 98 | 3 | 0 | 0 | 0 | < 0.001 | 1 | 0 | 0 | 0 | 100 | 101 | 0.065 |
| *etr-1(lq61) no Ex* | 79 | 14 | 6 | 0 | 1 |  | 1 | 1 | 0 | 4 | 94 | 100 |  |

\*compared to matched *etr-1(lq61) no Ex* at AQR position 1 and PQR position 5.

To further test the requirement of *etr-1* function in muscle, we used cell-specific genome editing involving transgenic expression of synthetic guide (sg)RNAs and Cas9. *The etr-1-* specific sgRNA (Figure 3D) was expressed from the ubiquitous U6 promoter, and Cas9 expression was driven from the body wall muscle-specific *myo-3* promoter (Dickinson *et al*. 2013). Animals harboring the muscle-specific genome editing transgene displayed in AQR/PQR directional defects similar to *lq61* and *lq133* mutations (Table 1).

### CRISPR-seq, a next-generation sequencing technique to detect cell-specific genome edits

Genome edits in mixed genotypic samples are generally detected by the T7E1 assay (Cong *et al*. 2013), which uses amplicon sequencing and digestion with T7E1 nuclease which cleaves at areas of heteroduplex DNA, such as formed by one strand of wild-type DNA forming a heteroduplex with an edited strand. The T7E1 assay was not successful at detecting genome edits in muscle-specific *etr-1[CRISPR)* strains that showed AQR/PQR defects (data not shown). At the time of Q migrations, body wall muscle cells represent approximately 10% of the cells of the L1 larva, raising the possibility that the T7E1 assay lacked the resolution to detect genome edits at this ratio.

We developed a next-generation sequencing protocol to detect cell-specific genome edits that we call CRISPR-seq (see Methods for a detailed protocol). First, genomic DNA was isolated from L3 animals harboring the cell-specific CRISPR transgene. L4 and adult animals were avoided because of the proliferation of germ line cells in these animals. Next, two-step amplicon PCR was used with gene-specific primers flanking the predicted edit site, which also incorporated the Illumina clustering and sequencing primers and indices, resulting in a standard Illumina amplicon sequencing library. These CRISPR-seq libraries were then sequenced on the Illumina platform, and reads aligned to the genome. The Integrated Genome Viewer (IGV) was used to determine the absence rate of single nucleotides among the reads aligning to the region. The absence rates were plotted versus genomic position, resulting in the graph in Figure 4A. *etr-1* CRIPR-seq resulted in a high frequency of missing bases near the PAM site, with a graded reduction toward the 3’ end. Of note, only a subset of reads showed deleted bases (4% or less), but significantly more than CRISPR-seq on wild-type animals (Figure 5A), which showed few or no deletions. In L3 animals, approximately 10% of the cells are body wall muscle, but we observed 4% or fewer reads with deletions. If deletions remove the primer sites, they will not be amplified and included in the library. Furthermore, some deletions that do amplify might not align due to their size or position relative to the amplicon primers. Thus, CRISPR-seq was not a quantitative measure of cell-specific genome editing, but did demonstrate that genome edits occurred in cell-specific conditions (i.e. mixed genotypic populations of muscle cells and other cells).

**Figure 5.**
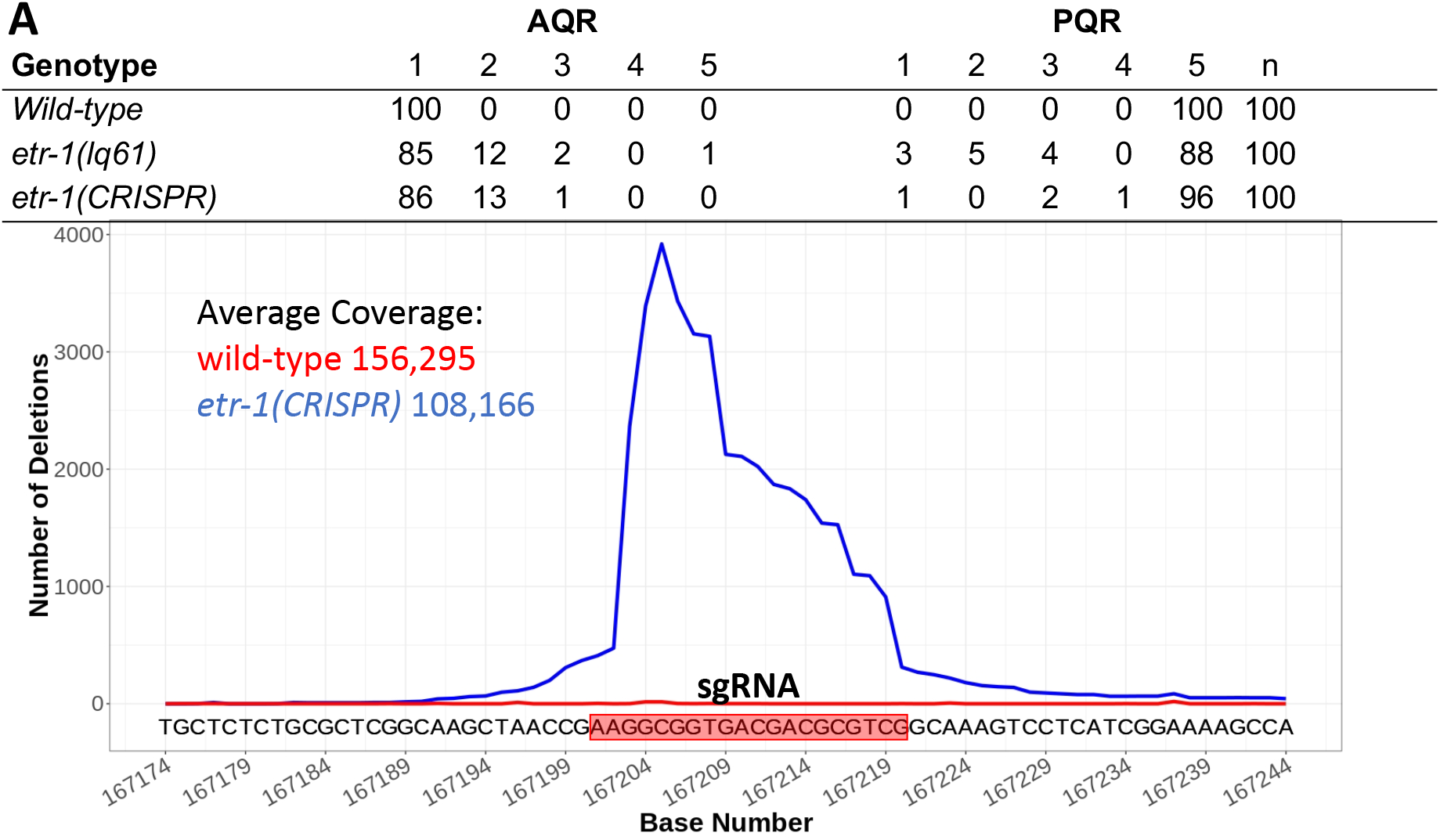
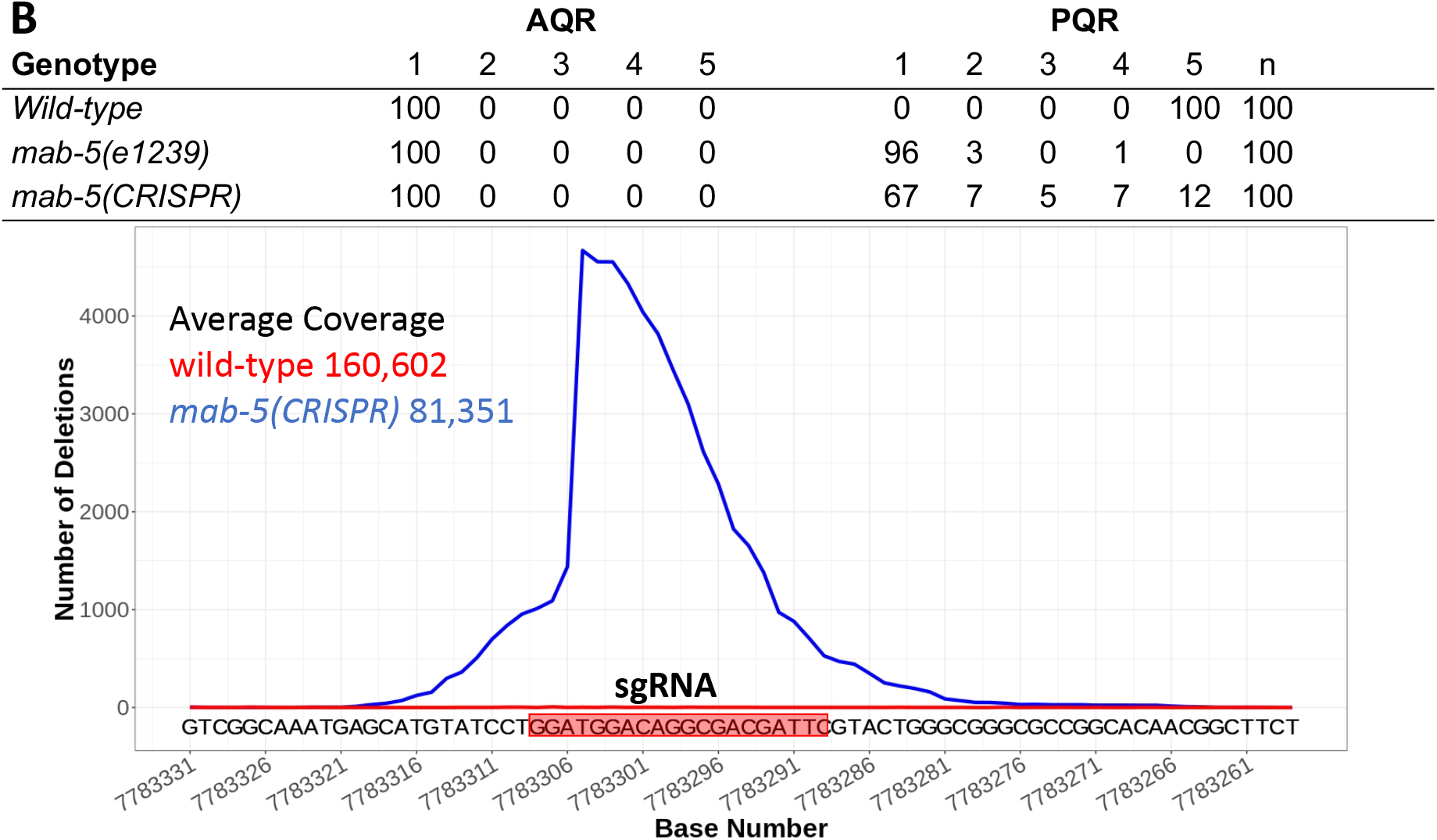
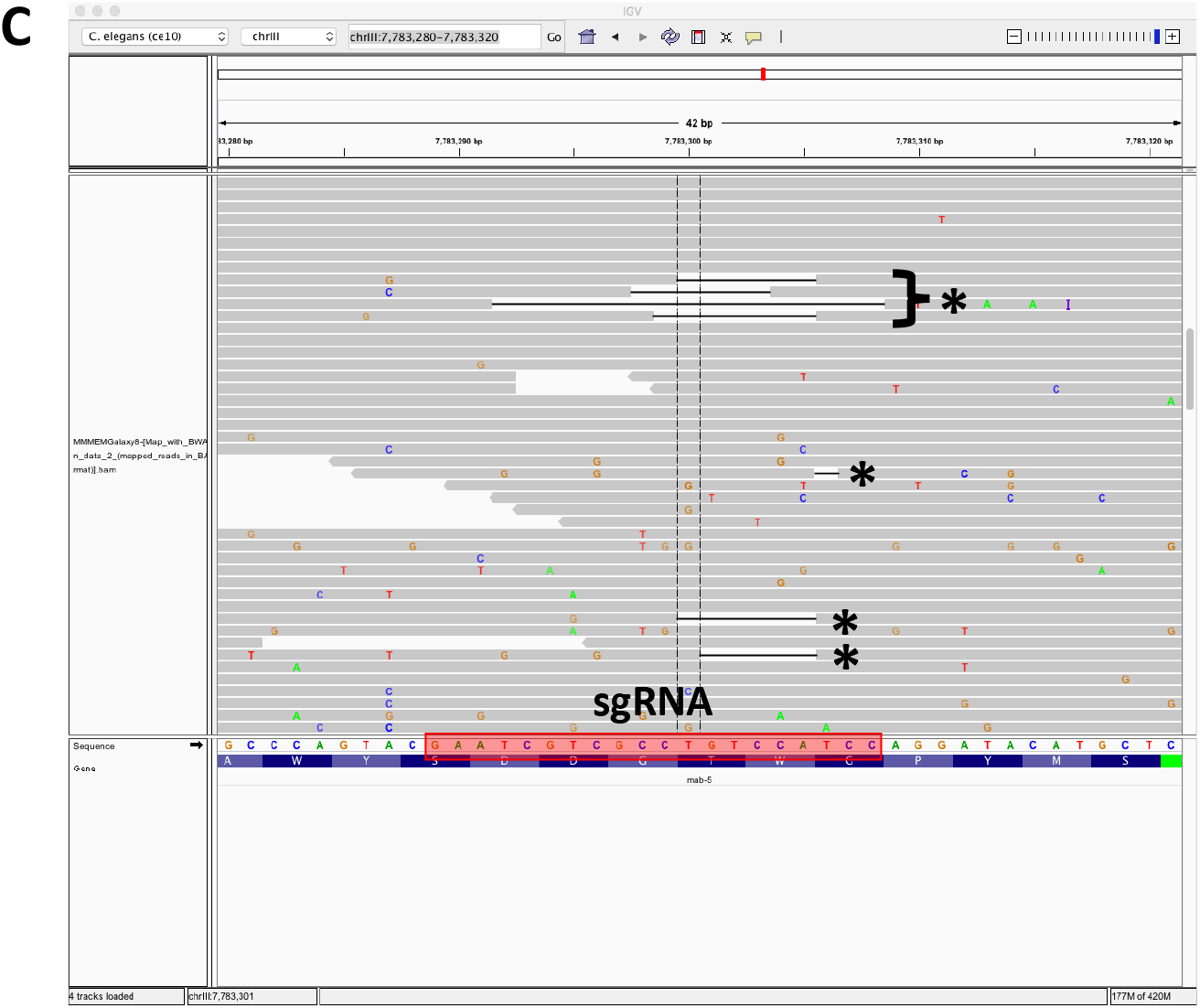
CRISPR-seq of *etr-1* and *mab-5* cell-specific genome editing. A) The table shows AQR and PQR migration in wild-type, an *etr-1(lq61)* mutant, and a transgenic *etr-1(CRISPR)* animals with ubiquitous *etr-1* sgRNA expression and Cas9 expression from the *myo-3* promoter expressed in body wall muscle. The graph shows the results of an amplicon sequencing experiment, with the chromosome position in nucleotides (X-axis) and the number of deletions at each position in wild-type (red) and *etr-1(CRISPR)* (blue) (Y-axis). The sgRNA sequence used is indicated in the red box. The average coverage in amplicon sequencing experiment across the region is indicated. B) A CRISPR-seq experiment on cell-specific *mab-5* CRISPR, as described for *etr-1* n (A). The *mab-5* sgRNA was expressed ubiquitously, and Cas9 was expressed from the Q-cell-specific *egl-17* promoter. C) A screenshot from Integrated Genome Viewer showing alignment of *mab-5* CRISPR-seq reads. Reads with deletions are indicated with asterisks. The sgRNA sequence is in the red box, and is the reverse complementary strand to the site depicted in Figure 5B.

As proof-of-principle, we conducted cell-specific genome editing and CRISPR-seq on another locus, the *mab-5* gene (Figure 5B). *mab-5* loss-of-function results in nearly completely penetrant PQR anterior migration. We expressed Cas9 from the Q-cell-specific *egl-17* promoter, which resulted in 88% defects in PQR migration, with 67% migrating fully anteriorly to the normal position of AQR (Figure 5B). CRISPR-seq identified a similar pattern of genome editing around the sgRNA site, with high frequency near the PAM site and a gradual decline to the 3’ of the site (Figure 5B). *egl-17* is expressed in the two Q cells, their descendants, and two cells in the head, representing one percent or fewer of the cells in L3/L4 animals. This indicates that CRISPR-seq has resolution to detect low-frequency genome editing events. In sum, CRISPR-seq is an effective method to detect low-frequency genome edits such as those that occur in cell-specific genome editing in multicellular animals.

### *etr-1* interacts with *wnt* mutations

Results presented here indicate that *etr-1* does not affect early Q neuroblast protrusion and migration, but does affect Q descendant migration, including direction of Q descendant migration. The five *C. elegans* Wnts have multiple roles in Q descendant migration. EGL-20 is required for expression of MAB-5 in QL, via canonical Wnt signaling involving BAR-1/ß-catenin to drive posterior migration. In *egl-20* mutants, PQR migrates posteriorly to the head (Table 3). EGL-20 and the other four Wnts CWN-1, CWN-2, LIN-44, and MOM-2 also have redundant roles in later Q descendant migration (Zinovyeva *et al*. 2008; Josephson *et al*. 2016), with double mutants resulting in failures of migration in the anterior-posterior axis, as well as directional defects (Table 3). For example, in *egl-20* double mutants with *cwn-1, cwn-2*, and *lin-44*, both AQR and PQR fail in their anterior migrations at a significantly higher frequency than additive effects of single mutants alone; and *cwn-1; cwn-2* double mutants have synergistic failures in AQR anterior migration (Table 3). *etr-1; egl-20* mutants display synergistic failures of anterior PQR migration in *egl-20; etr-1* double mutants, similar to *wnt* mutations. *etr-1; cwn-2* mutants also display synergistic AQR anterior migration defects (Table 3). Failures in anterior AQR and PQR migration are yet more severe in *etr-1; egl-20 cwn-2* triple mutants. These data indicate that *etr-1* modifies the *egl-20* phenotype in a manner similar to *wnt* mutations, and might act with *wnts* or in parallel to guide Q descendant migrations in the A-P axis.

**Table 3.** *etr-1* interactions with *Wnt* mutations.

| Genotype | AQR |  |  |  |  | *p* | PQR |  |  |  |  | *n* | *p* |
| --- | --- | --- | --- | --- | --- | --- | --- | --- | --- | --- | --- | --- | --- |
|  | 1 | 2 | 3 | 4 | 5 |  | 1 | 2 | 3 | 4 | 5 |  |  |
| *etr-1(lq61)* | 85 | 12 | 2 | 0 | 1 |  | 3 | 5 | 4 | 0 | 88 | 100 |  |
| *egl-20(gk453010)* | 99 | 1 | 0 | 0 | 0 |  | 94 | 5 | 0 | 1 | 0 | 100 |  |
| *egl-20(n585)* | 100 | 0 | 0 | 0 | 0 |  | 97 | 2 | 1 | 0 | 0 | 100 |  |
| *egl-20(mu39)* | 94 | 6 | 0 | 0 | 0 |  | 36 | 7 | 2 | 3 | 52 | 100 | <sup>1</sup><0.001 |
| *cwn-1(ok546)* | 71 | 23 | 6 | 0 | 0 |  | 0 | 0 | 0 | 0 | 100 | 100 |  |
| *cwn-2(ok895)* | 91 | 9 | 0 | 0 | 0 |  | 0 | 0 | 0 | 4 | 96 | 100 |  |
| *lin-44(e1792)* | 99 | 1 | 0 | 0 | 0 |  | 0 | 0 | 0 | 0 | 100 | 100 |  |
| *cwn-1(ok546); egl-20(n585)* | 5 | 23 | 35 | 34 | 3 | <sup>2</sup><0.001 | 0 | 12 | 34 | 51 | 2 | 100 | <sup>1</sup><0.001 |
| *cwn-1(ok546); cwn-2(ok895)* | 36 | 57 | 6 | 1 | 0 | <sup>3</sup><0.001 | 0 | 0 | 0 | 0 | 100 | 100 |  |
| *lin-44(n1792); egl-20(gk453010)* | 88 | 12 | 0 | 0 | 0 |  | 61 | 30 | 7 | 0 | 2 | 100 | <sup>1</sup><0.001 |
| *egl-20(n585) cwn-2(ok895)* | 16 | 67 | 17 | 0 | 0 |  | 13 | 74 | 13 | 0 | 0 | 100 |  |
| *egl-20(n585) cwn-2(ok895); cwn-1(ok546)* | 21 | 70 | 8 | 1 | 0 |  | 24 | 54 | 21 | 1 | 0 | 100 |  |
| *etr-1(lq61); egl-20(gk453010)* | 87 | 11 | 1 | 1 | 0 |  | 71 | 20 | 8 | 0 | 1 | 100 | <sup>1</sup><0.001 |
| *etr-1(lq61); egl-20(mu39)* | 69 | 29 | 9 | 1 | 0 |  | 4 | 4 | 3 | 12 | 77 | 100 | <sup>4</sup><0.001 |
| *etr-1(lq61); cwn-2(ok895)* | 56 | 35 | 7 | 0 | 2 | <sup>5</sup>0.003 | 3 | 1 | 2 | 11 | 83 | 100 |  |
| *etr-1(lq61); lin-44(e1792)* | 81 | 16 | 3 | 0 | 0 |  | 0 | 1 | 1 | 3 | 95 | 100 |  |
| *etr-1(lq61); egl-20(n585) cwn-2(ok895)* | 41 | 40 | 15 | 4 | 0 | <sup>6</sup><0.001 | 43 | 34 | 13 | 10 | 0 | 100 | <sup>6</sup><0.001 |

<sup>1</sup>compared to *egl-20(gk453010)* and *egl-20(n585)* at PQR position 1.
<sup>2</sup>compared to *cwn-1(ok546)* at AQR position 1.
<sup>3</sup>compared to the additive effects of *cwn-1(ok546)* and *cwn-2(ok895)* at AQR position 1.
<sup>4</sup>compared to *egl-20(mu39)* at PQR position 1.

## Discussion

### ETR-1/CELF1 isoforms containing exon 8 are required for neuronal migration

CELF1 molecules are RNA-binding proteins that mediate multiple aspects of RNA processing, including alternative splicing (reviewed in(Dasgupta and Ladd 2012)). A forward genetic screen for mutations affecting the migration of the Q neuroblast descendant neurons AQR and PQR identified the *lq61* mutation in the *etr-1* gene, which encodes a molecule similar to mammalian CELF1 (Table 1 and Figures 2 and 3). *etr-1* was previously shown to be expressed in body wall muscle and to control muscle differentiation (Milne and Hodgkin 1999). RNAi of *etr-1* resulted in embryonic lethality at the paralyzed, arrested at two-fold stage (Pat) phenotype indicative of severe body wall muscle defects. The *etr-1(lq61)* allele isolated here was viable and fertile and did not show the Pat phenotype. *lq61* caused a premature stop codon in exon 8 of *etr-1*, an exon that displays extensive alternative splicing (WormBase web site, http://www.wormbase.org, release WS274, 9/27/2019)(Boateng *et al*. 2017) (Figure 3). Thus, *etr-1(lq61)* likely affects a subset of *etr-1* transcripts containing exon 8. Indeed, transgenic expression of *etr-1(+)* genomic DNA rescued *etr-1(lq61)* AQR/PQR migration defects, and CRISPR/Cas9 mediated induction of an independent mutation in exon 8 recapitulated the *etr-1(lq61)* phenotype (Table 1 and Figure 3).

ETR-1 transcripts containing exon 8 are required for AQR/PQR neuronal migration. Exon 8 encodes for a poly-glutamine region between the second and third RNA. Recognition Motifs (RRMs) and does not encode any of the three RRMs in ETR-1 (Figure 3). In other RNA binding proteins, poly-glutamine regions have been shown to modulate interaction with other splicing proteins and to affect target RNA splicing (Singh *et al*. 2011). Thus, ETR-1 isoforms containing exon 8 might affect a subset of normal RNA targets, which are disrupted by *etr-1(lq61)*. This might explain why complete knockdown of *etr-1* results in embryonic lethality and the Pat phenotype, whereas *etr-1(lq61)* results in viable and fertile animals with AQR/PQR migration defects.

### ETR-1 is required in body wall muscle for AQR/PQR neuronal migration

Previous work showed that *etr-1* is expressed strongly in body wall muscles (Milne and Hodgkin 1999). Recently, *etr-1* expression was described in most if not all cells of the embryo (Boateng *et al*. 2017). Two lines of evidence suggest that ETR-1 is required in the body wall muscles for proper AQR/PQR migration. First, expression of *etr-1(+)* coding region in Q cells using the *egl-17* promoter did not rescue AQR/PQR defects of *etr-1(lq61)*, whereas expression in body wall muscle cells via the *myo-3* promoter did rescue (Table 2). Second, cell-specific CRISPR/Cas9 mediated knockout of *etr-1* exon 8 in the body wall muscle cells resulted AQR/PQR defects similar to *etr-1(lq61)* (Table 1). Despite expression in most if not all cells, *etr-1* isoforms with exon 8 were required in the body wall muscle for proper AQR/PQR migration *(e.g*. non-autonomously). It is possible that *etr-1* isoforms have functions in other cells, including the Q cells, not detected here. Indeed, *etr-1* controls multiple aspects of germ cell development including oocyte maturation and germ cell engulfment after programmed cell death (Boateng *et al*. 2017). Possibly, different ETR-1 isoforms are expressed in distinct tissues and mediate distinct functions. For example, expression of isoforms with exon 8 might be restricted to body wall muscles. Future experiments will address these questions.

### *etr-1(lq61)* interacts with mutations in Wnt signaling

Q neuroblast migration occurs in two phases (Figure 1). After hatching, initial Q cell migration involves extension of protrusions to the anterior and posterior for QR and QL respectively, with subsequent migration of the cell bodies. despite extensive testing, no role for Wnt signaling has been identified in this first phase (Josephson *et al*. 2016), which rather involves the transmembrane receptors PTP-3/LAR and UNC-40/DCC, and Fat-like Cadherins CDH-3 and CDH-4 (Honigberg and Kenyon 2000; Middelkoop *et al*. 2012; Sundararajan and Lundquist 2012; Sundararajan *et al*. 2014; Ebbing *et al*. 2019). *etr-1(lq61)* mutants showed no defects in initial Q protrusion or migration (Figure 4), suggesting that ETR-1 exon 8 isoforms are not involved in initial migration.

The second phase of Q descendant migration involves both canonical and non-canonical Wnt signaling (see (Eisenmann 2005; Zinovyeva *et al*. 2008; Ji *et al*. 2013; Middelkoop and Korswagen 2014; Josephson *et al*. 2016)). EGL-20/Wnt activates expression of MAB-5/Hox in QL descendants which drives continued posterior migration, and expression of all five Wnt ligands in distinct regions of the anterior-posterior axis controls Q descendant migration, including AQR and PQR, via non-canonical pathways not involving BAR-1/ß-catenin (Chapman *et al*. 2008; Zinovyeva *et al*. 2008; Harterink *et al*. 2011; Josephson *et al*. 2016). Single, double, and triple *Wnt* mutation combinations reveal redundant functions in AQR/PQR migration, including misdirection (canonical) and failures to migrate completely (non-canonical), similar to *etr-1(lq61)* (Table 3). Double mutants of *Wnt* mutations with *etr-1(lq61)* revealed redundant functions in AQR/PQR migration. In other words, *etr-1(lq61)* acts redundantly with *Wnt* single mutants in a manner similar to *Wnt* double mutants. These data are consistent with ETR-1 exon 8 isoforms acting with Wnt signaling, or in a pathway parallel to Wnt signaling, to control AQR/PQR migration.

Wnt ligands, including EGL-20, CWN-1, CWN-2, and MOM-2, are expressed in body wall muscle cells and other tissues (Gleason *et al*. 2006; Pan *et al*. 2006; Harterink *et al*. 2011). As ETR-1 exon 8 isoforms act non-autonomously in the body wall muscle cells to control AQR/PQR neuronal migration, they might regulate the production of a signal from body wall muscles that controls AQR/PQR migration. This could be Wnt production itself, or a signal that acts in parallel to Wnt signaling to control AQR/PQR migration. Future experiments will test these ideas.

### CRISPR-seq, a new method to detect rare genome editing events *in vivo*

Cell-specific and conditional expression of Cas9 resulting in cell-specific or conditional genome editing is a powerful tool for analysis of cell- and tissue-specific effects of mutations (Shen *et al*. 2014). The T7E1 assay is used to detect rare genome editing events in mixed populations of cells or organisms. Cell-specific genome editing of *etr-1* in body wall muscles and *mab-5* in Q cells resulted in AQR/PQR migration defects similar to the mutations alone (Table 1 and Figure 5). However, the T7E1 assay did not detect genome edits in these mosaic animals (data not shown). We used amplicon sequencing to detect rare genome editing events in these animals (CRISPR-seq). Primers were designed, flanking the sgRNA site, for amplification in a two-step process that adds the Illumina clustering and sequencing primer sites to the amplicon library (see materials and Methods). The libraries were then sequenced using the Illumina Miseq platform, and reads were aligned to the genome. Base coverage at each site in the amplicon was determined using the Integrated Genome Viewer, and was plotted as base position against number of missing bases in the reads (Figure 5).

These data show that rare genome editing events were readily detected. At the time of DNA extraction, ~10% of all cells were body wall muscle cells, yet fewer than 4% of reads displayed a deletion. Therefore, it is likely that not all genome editing events are detected by this method. For example, events that remove the amplicon primer sites or events that result in very short reads that do not align, would not be detected. It is also possible that genome editing is not completely efficient, which would also lower read counts. For the *mab-5* Q cell experiment, the Q cells represented <1% of all cells when DNA was harvested. However, *egl-17* expression begins in many other cells besides the Q cells later in larval development (e.g. the P cells) (Burdine *et al*. 1998). CRIPR-seq is not a quantitative method for cell-specific genome editing, but readily detects rare genome editing events in cell-specific genome editing experiments *in vivo*.

### Summary

In this work we describe a novel role of the CELF1-family RNA-binding protein in neuroblast migration. Surprisingly, this is a non-autonomous role, as ETR-1 was required in the body wall muscle cells for neuroblast migration. We speculate that ETR-1 is involved in the production of a signal from the body wall muscles that provides guidance and migration information to the Q neuroblasts as they migrate in the anterior-posterior axis. Interactions with Wnt signaling suggest ETR-1 could be acting with Wnt signaling, possible in the production of the Wnt signal from body wall muscles, or with an as-yet unidentified signaling system in parallel to Wnt. The *etr-1* mutations described here affect only ETR-1 isoforms that contain exon 8, which encodes a polyglutamine-rich region, which can interact with other splicing factors and mediate target-specific interaction. Thus, *etr-1(lq61)* might affect only a subset of the normal RNA targets of ETR-1 processing. Finally, we describe a method, CRISPR-seq, that utilize amplicon sequencing to detect rare genome editing events in genetically mosaic animals and is an alternative to the T7E1 assay. In humans, CELF 1 disruption is involved in a spectrum of neuromuscular and other diseases, consistent with a role of ETR-1 in muscle development and function. Our results suggest that cell or neuronal migration might be an aspect of some of these disorders.

## Acknowledgments

The authors thank E. Struckhoff for technical assistance, and the members of the Ackley and Lundquist labs for discussion. This work was funded by NIH grant R21NS100483 to E.A.L., and the Kansas Infrastructure Network of Biomedical Research Excellence NIH grant P20GM103418. Some strains were provided by the CGC, which is funded by NIH Office of Research Infrastructure Programs (P400D010440). Next generation Illumina sequencing was done at the KU Genome Sequencing Core laboratory supported by NIH grant P20GM103638.

